# Determine independent gut microbiota-diseases association by eliminating the effects of human lifestyle factors

**DOI:** 10.1101/2021.01.14.426764

**Authors:** Xin Wang, Yuqing Yang, Jianchu Li, Rui Jiang, Ting Chen, Congmin Zhu

## Abstract

Human lifestyle and physiological variables on human disease risk have been revealed to be mediated by gut microbiota. Low concordance between many case-control studies for detecting disease-associated microbe existed and it is likely due to the limited sample size and the population-wide bias in human lifestyle and physiological variables. To infer association between whole gut microbiota and diseases accurately, we propose to build machine learning models by including both human variables and gut microbiota based on the American Gut Project data, the largest known publicly available human gut bacterial microbiota dataset. When the model's performance with both gut microbiota and human variables is better than the model with just human variables, the independent association of gut microbiota with the disease will be confirmed. We found that gut microbes showed different association strengths with different diseases. Adding gut microbiota into human variables enhanced the association strengths with inflammatory bowel disease (IBD) and unhealthy status; showed no effect on association strengths with Diabetes and IBS; reduced the association strengths with small intestinal bacterial overgrowth, *C. difficile* infection, lactose intolerance, cardiovascular disease and mental disorders. Our results suggested that although gut microbiota was reported to be associated with many diseases, a considerable proportion of these associations may be spurious. We also proposed a list of microbes as biomarkers to classify IBD and unhealthy status, and validated them by reference to previously published research.

**IMPORTANCE:** we reexamined the association between gut microbiota and multiple diseases via machine learning models on a large-scale dataset, and by considering the effect of human variables ignored by previous studies, truly independent microbiota-disease associations were estimated. We found gut microbiota is associated independently with IBD and overall health of human, but more evidence is needed to judge associations between microbiota and other diseases. Further functional investigations of our reported disease-related microbes will improve understanding of the molecular mechanism of human diseases.

## INTRODUCTION

The human intestines are home to a dense microbial community, collectively known as the gut microbiota (1). The gut microbiota forms a complex ecosystem and performs a wide range of functions with far-reaching impacts on human health, including extracting energy from the digestive system, preventing colonization by pathogens, promoting immune homeostasis, producing important metabolites, and even communicating with the central nervous system via the gut-brain axis (2). So, it is thought to play an important role in the development of many diseases, including inflammatory bowel disease (3), *Clostridium difficile* infection (4), Diabetes (5), cardiovascular disease (6), and mental health disorders (7). The gut microbiota determines certain host characteristics and responds to host variables, such as human lifestyle and physiological variables, which can be reflected in the microbial composition (8). Therefore, a considerable part of the human variables on human health and disease risk may be mediated or modified by gut microbiota.

With the development of high-throughput sequencing technology, we are now able to sequence the hypervariable regions of the 16S rRNA gene and cluster into operational taxonomic units (OTUs) to profile the taxonomic composition of the microbial community in an environmental sample (9). Over the last few years, many case-control studies have been conducted to collect microbial 16S rRNA gene datasets from human fecal samples to explore the associations among the gut microbial community and human diseases to reveal disease-specific microbial biomarkers (10, 11). However, many investigations showed low concordance on the discovered disease-associated microbes and one obvious example is studies about obesity. Gut microbiota reported by multiple studies of which abundance is differential between obese and lean individuals is inconsistent (12). Furthermore, M. A. Sze and P. D. Schloss (13) comprehensively analyzed the results of several obesity-related studies. They found that the statistical detection power of a small-sample study was insufficient, and the ratio of abundance of *Bacteroidetes* and *Firmicutes* was not associated with obesity. In addition, a recent construction of a large dataset from the Swedish population did not reveal an apparent microbial signature associated with irritable bowel syndrome (IBS) as previously reported in the literature, and the heterogeneity of the microbial community among IBS patients was higher than that among healthy individuals (14).

There are two possible reasons which may result in low concordance in previous studies. One is the limitation of the sample size. Generally, there are thousands of microorganisms with a wide range of abundance levels in intestinal samples. Due to the high cost of building a large-scale dataset consisting of both gut microbiota information and elaborate human variables (15), researchers can only afford to sequence dozens or hundreds of samples to explore disease-associated microbes via statistical models. Thus, the model overfitting is common and thus reduces the reliability of the inferred results. The other critical shortcoming is neglecting the influence of host variables, which makes it difficult for researchers to confirm whether the calculated gut microbial-disease associations indicate the true interactions between microbes and the progression of diseases. The alternative possibility is that microbes are only related to certain host variables, and as a result, they are associated indirectly with diseases(16).

Therefore, a large-scale dataset containing information on both gut microbial community and host variables is required for accurate identification of microbiota-disease associations. Fortunately, the American Gut Project (AGP), which comprised thousands of 16s rRNA gene sequencing samples and a rich human variables set related to human lifestyle and physiological variables and diseases, has been carried out worldwide (17). Today, the AGP has sequenced more than 15,000 samples, which significantly expands human gut microbiota’s existing data. Most importantly, it provides a rich resource for each sample with information on gut microbiota, human lifestyle factors, and diseases. The goal of this study is to explore the relationship between these entities using this dataset.

Our approach is different from traditional association inference analysis, which tries to estimate the relationship between a single microbe and a disease. We focus on determining whether the whole gut microbiota is independently associated with human diseases by eliminating the influence of lifestyle factors using machine learning methods (ML). The strength of association between gut microbiota and disease is evaluated by the classification performance of the ML models built with the microbiota. Although researchers have built large microbial datasets by merging different studies to explore the effect of gut microbiota on predicting diseases and mortality risks (18–20) via ML approaches, they neglected the impact of human lifestyle factors, resultant in the magnified predictive power of the microbiota. It is because human lifestyle factors influence both the gut microbiota and the disease progression. Besides, human lifestyle factors between enrolled healthy individuals (controls) and patients (cases) can be significantly different, and such differences could become the main contributor to the predictive power of the disease. It is not surprising to build a well-performed disease classification model using both gut microbiota and human lifestyle and physiological variables in this condition. However, we argue that the independent gut microbiota-disease associations are real, only when the models’ classification performance with both gut microbiota and human variables is significantly better than the model built with just human lifestyle factors (Fig. 1). Conversely, when the models’ classification performance is inferior to the human-lifestyle-built model, either gut microbiota may not be associated with diseases, or a more suitable data enrollment criterion is needed.

**Fig.1.**
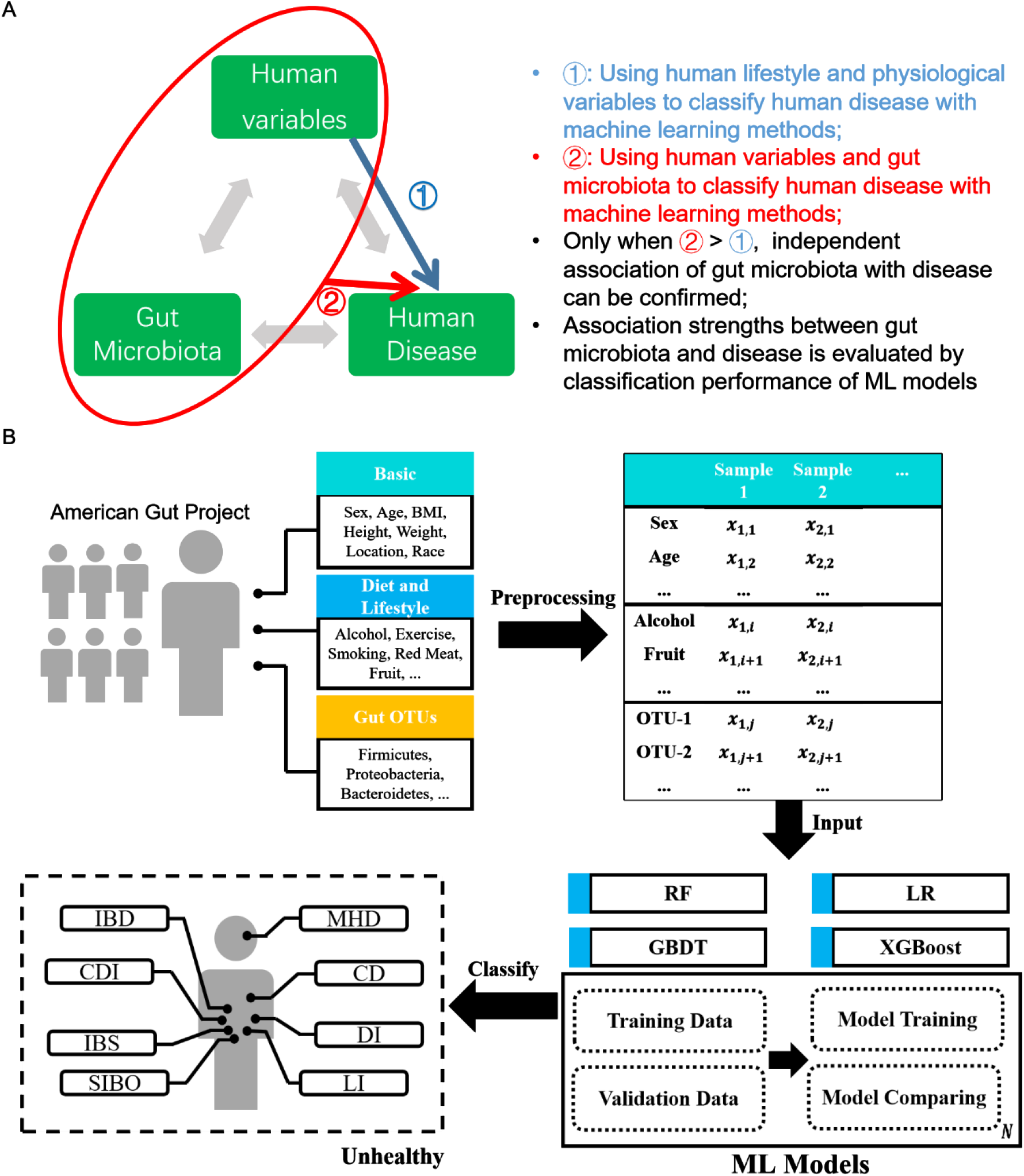
Workflow of disease classification models construction. A) We propose to build association models by including both human variables and gut microbiota from American Gut Project, and assumed that when the performance of the model with both gut microbiota and human variables is better than the model with just human variables, the independent association of gut microbiota with the disease can be confirmed. B) We classified eight diseases (IBD: Inflammatory Bowel Disease; CDI: C. Difficile Infection; IBS: Irritable Bowel Syndrome; SIBO: Small Intestinal Bacterial Overgrowth; DI: Diabetes; LI: Lactose Intolerance; CD: Cardiovascular Disease; MD: Mental Disorder) with human variables (physiological characteristics, lifestyle, location and diet) and gut microbial community data (OTUs) obtained from the American Gut Project (AGP) database using four machine learning techniques (Random Forest, Gradient Boosting Decision Tree, Logistic Regression and eXtreme Gradient Boosting).

Following the argument, we explored the classification power of the gut microbiota and human variables on multiple diseases using the AGP data with ML classification models. Key OTUs and human variables, consisting of lifestyle and dietary factors, were identified with high validity using multiple machine learning methods (ML). The performance of OTUs and human variables was compared comprehensively to show the difference between their contributions to diseases. In addition, considering the widespread associations of gut microbes with multiple diseases, we use OTUs to judge the overall health status of human, and individuals with at least one disease were classified as the unhealthy. Although lots of associations with diseases were identified previously, our results showed that adding gut microbiota into human variables only enhanced the association strengths with inflammatory bowel disease (IBD) and unhealthy status. In addition, we reported the top 10 features (OTUs or phenotypes) used in the classification of these diseases, most of which were supported by previously published studies.

## RESULTS

### Characteristics of the dataset

Dataset used in this study consisted of 7,571 samples with 517 OTUs and 30 human variables (See MATERIALS AND METHODS for details). For human variables, there were 6 variables related to individuals’ physiological characteristics (age, sex, height, weight, body mass index (BMI) and race), 2 related to lifestyle choices (exercise and smoking frequencies), 3 related to location (latitude, elevation and country), and 19 related to diet (frequencies of fruit, high-fat red meat, alcohol, and so on). For every sample, labels of eight diseases [cardiovascular disease (CD), small intestinal bacterial overgrowth (SIBO), mental disorders (MD), lactose intolerance (LI), diabetes (DI), IBD, irritable bowel syndrome (IBS), *C. difficile* infection (CDI) and Diabetes (DI)] that have been reported to be related to gut microbiota were extracted. Besides, a disease label named ‘unhealthy (UH)’ was added if a sample had at least one of eight diseases. The characteristics of the dataset, the demographic details of samples and the number of male and female patients for each disease are shown in Table S1 and S2.

For every disease, ML classification models were constructed using five types of features respectively: human variables only (Meta), OTU abundance only (OTUab), OTU occurrence only (OTUoc), both human variable data and OTU abundance (Meta-OTUab), and both human variable data and OTU occurrence (Meta-OTUoc). Models were trained and compared on identical training and validation data. For each type of feature, the best model was selected according to the AUC score. By comparing the performance of models with only Meta and models with both human variables and OTU information (Meta-OTUab and Meta-OTUoc), all diseases were classified into three categories: adding gut microbiota a) could improve, b) didn’t affect or c) reduced disease classification performance.

Adding gut microbiota into human variables enhanced the association strengths with IBDAs a global public health concern, the incidence and prevalence of IBD, which is caused by gut dysbiosis, is increasing in developed and developing countries (21, 22). In this study, 413 IBD patients from the AGP were included in the final dataset, and the results of the best models using five types of features are shown in Table 1. The model using human variable data only (Meta) as the feature achieved an AUC of 0.72680±0.00874. Interestingly, the AUCs of using OTUab alone (0.71404±0.01229) and OTUoc alone (0.70693±0.01536) did not differ significantly from that obtained using Meta (*P* = 0.0096), indicating that gut microbiota alone is as good a classifier of IBD as human variables. Additionally, the AUCs obtained using the combination of Meta and OTUs (Meta-OTUab: 0.74706±0.01421 and Meta-OTUoc: 0.75464±0.01855) were significantly higher than those obtained using Meta alone (*P* = 0.00037 and *P* = 0.00057, respectively), suggesting that adding gut microbiota into human variables significantly enhanced the association strengths with IBD (Table 1) and that the independent association of gut microbiota with IBD can be confirmed. It is noteworthy that the Meta-OTUoc achieved higher AUCs than those achieved with Meta-OTUab, which implied that, compared to the abundance of gut microbes, their occurrences are better features for the classification of IBD.

**Table 1.** Comparing AUC values of nine diseases using five feature types.

| Feature type |  | Meta | OTUab | OTUoc | Meta-OT<br>Uab | Meta-OT<br>Uoc |
| --- | --- | --- | --- | --- | --- | --- |
| IBD | AUC | 0.72680 ± 0.00874 | 0.71404 ± 0.01229 | 0.70693 ± 0.01536 | 0.74706 ± 0.01421 | 0.75464 ± 0.01855 |
|  | p-value | - | 0.00965 | 0.00256 | 0.00037 | 0.00057 |
| DI | AUC | 0.84658 ± 0.02231 | 0.66839 ± 0.02136 | 0.66216 ± 0.03878 | 0.82068 ± 0.02053 | 0.83591 ± 0.01429 |
|  | p-value | - | <0.0001 | <0.0001 | 0.05067 | 0.0864 |
| IBS | AUC | 0.68448 ± 0.00780 | 0.59820 ± 0.00995 | 0.56972 ± 0.01460 | 0.66473 ± 0.01118 | 0.66950 ± 0.01386 |
|  | p-value | - | <0.0001 | <0.0001 | 0.00107 | 0.00967 |
| SIBO | AUC | 0.74413 ± 0.0234 | 0.60070 ± 0.02934 | 0.62955 ± 0.01989 | 0.67722 ± 0.02206 | 0.69628 ± 0.02600 |
|  | p-value | - | <0.0001 | <0.0001 | 0.00001 | 0.00003 |
| CDI | AUC | 0.84091 ± 0.01890 | 0.73271 ± 0.01730 | 0.68482 ± 0.01535 | 0.76280 ± 0.01307 | 0.75288 ± 0.03125 |
|  | p-value | - | <0.0001 | <0.0001 | 0.00001 | 0.00015 |
| LI | AUC | 0.81647 ± 0.00751 | 0.52619 ± 0.02473 | 0.50884 ± 0.02027 | 0.77253 ± 0.00730 | 0.79129 ± 0.00749 |
|  | p-value | - | <0.0001 | <0.0001 | <0.0001 | <0.0001 |
| CD | AUC | 0.83432 ± 0.01255 | 0.53990 ± 0.02785 | 0.57160 ± 0.02203 | 0.70787 ± 0.02003 | 0.73538 ± 0.02834 |
|  | p-value | - | <0.0001 | <0.0001 | <0.0001 | <0.0001 |
| MD | AUC | 0.62984 ± 0.01240 | 0.57123 ± 0.01472 | 0.55361 ± 0.01702 | 0.58390 ± 0.01853 | 0.58468 ± 0.01997 |
|  | p-value | - | 0.00005 | <0.0001 | 0.00006 | 0.0002 |
| UH | AUC | 0.70936 ± 0.00442 | 0.71976 ± 0.00533 | 0.71829 ± 0.00730 | 0.75626 ± 0.00859 | 0.76266 ± 0.00520 |
|  | p-value | - | 0.00016 | 0.02248 | <0.0001 | <0.0001 |

The model training process was repeated 10 times to assess the significance of differences in the model performance. P values were calculated by comparing with AUC scores of models using Meta respectively. The AUC scores were presented as the mean ± standard deviation.

Next, we assessed the relative roles of Meta or OTUs in the best ML model for classifying IBD with Meta, Meta-OTUab, and Meta-OTUoc. We ranked the features according to their weights and calculated the average rank after repeating the model training process ten times. As shown in Table 2, we found that the top 10 features for three types of features were distinct. For the model using human variable data only (Meta) as features, the top 10 most important human variables for classifying IBD comprised six dietary characteristics (the frequencies of vitamin B, alcohol, vitamin D, whole grain, probiotic, and red meat intake) and four basic physical characteristics (BMI, age, height, and weight), without geographical location features. For the model using Meta-OTUab as features, except for three dietary characteristics (the frequencies of vitamin B, vitamin D and probiotic intake), the other seven of the top 10 features for classifying IBD were all OTUs (four *Clostridiales*, one *Bacteroidales*, one *Erysipelotrichales* and one *Pasteurellales*). When using Meta-OTUoc as features to classify IBD, the results changed in that, except for two physical characteristics (age and height) and one OTU of *Enterobacteriales*, the other seven of the top 10 features were all OTUs of *Clostridiales*.

**Table 2.** Top ten most important features using three types of feature sets for IBD.

| Meta |  | Meta-OTUab |  | Meta-OTUoc |  |
| --- | --- | --- | --- | --- | --- |
| Feature Name | Rank | Feature Name | Rank | Feature Name | Rank |
| VITAMIN_B_SUPPLEMENT | 1.9 | VITAMIN_B_SUPPLEMENT | 1.2 | c_Clostridia;o_Clostridiales;f_Ruminococcaceae;g_Ruminococcus;s_ | 38.3 |
| ALCOHOL | 4.7 | c_Clostridia;o_Clostridiales;f_Lachnospiraceae;g_Coproccoccus;s_ | 5.1 | c_Clostridia;o_Clostridiales;f_Lachnospiraceae;g_;s_ | 43.2 |
| VITAMIN_D_SUPPLEMENT | 4.7 | c_Erysipelotrichi;o_Erysipelotrichales;f_Erysipelotrichaceae;g_Holdemania;s_ | 7.8 | c_Clostridia;o_Clostridiales;f_Ruminococcaceae;g_;s_ | 43.3 |
| BMI | 6.2 | VITAMIN_D_SUPPLEMENT | 12.2 | c_Clostridia;o_Clostridiales;f_Lachnospiraceae;g_Coproccoccus;s_ | 45.4 |
| AGE_CORRECTED | 8.2 | c_Clostridia;o_Clostridiales;f_Ruminococcaceae;g_;s_ | 17.8 | AGE_CORRECTED | 46.2 |
| HEIGHT_CM | 9.1 | PROBIOTIC | 19.5 | c_Clostridia;o_Clostridiales;f_;g_;s_ | 48 |
| WHOLE_GRAIN | 9.2 | c_Gammaproteobacteria;o_Pasteurellales;f_Pasteurellaceae;g_Haemophilus;s_parainfluenzae | 20.3 | c_Gammaproteobacteria;o_Enterobacteriales;f_Enterobacteriaceae;g_Klebsiella;s_ | 50.4 |
| PROBIOTIC | 10 | c_Bacteroidia;o_Bacteroidales;f_Bacteroidaceae;g_Bacteroides;s_ | 25.6 | HEIGHT_CM | 52.1 |
| WEIGHT_KG | 12 | c_Clostridia;o_Clostridiales;f_Ruminococcaceae;g_Ruminococcus;s_ | 28.7 | c_Clostridia;o_Clostridiales;f_Lachnospiraceae;g_[Ruminococcus];s_ | 54.4 |
| RED_MEAT | 12.2 | c_Clostridia;o_Clostridiales;f_Ruminococcaceae;g_Faecalibacterium;s_prausnitzii | 31.5 | c_Clostridia;o_Clostridiales;f_;g_;s_ | 56.6 |

### Adding gut microbiota showed no effect on association strengths with Diabetes and IBS

DI and IBS are widely reported to be closely related to gut microbes and some diet habits (3, 23); therefore, we hypothesized that adding microbes into human variables will improve the classification of these two conditions. As shown in Table 2, Meta provided the best performance in classifying both these two diseases. The maximal AUCs for classifying CDI with Meta were 0.84658±0.02231. The AUCs obtained using OTUab alone (0.66839±0.02136) and OTUoc alone (0.66216±0.03878) were significantly lower than those obtained using Meta. It is noteworthy that the AUCs obtained using the combination of Meta and OTUs (Meta-OTUab: 0.82068±0.02053 and Meta-OTUoc: 0.83591±0.01429) did not differ significantly from those obtained using Meta alone (*P* = 0.0506 and *P* = 0.0864, respectively), suggesting that adding gut microbiota into human variables showed no effect on association strengths with DI. Meta-OTUoc achieved higher AUCs than Meta-OTUab. The ROC curves of the models based on Meta-OTUab and Meta-OTUoc were close to those of the Meta-only based models (Fig. 2). According to the feature weights, we identified the top 10 most important human variables for classifying DI (Fig. 3 and Table S3). When using Meta as the feature set, the top 10 most important human variables for IBS comprised four basic physical characteristics (BMI, age, weight and height), two geographical location features (elevation and latitude), three dietary characteristics (the frequencies of alcohol, milk cheese and milk intake) and one lifestyle habit (exercise frequency). For the model with Meta-OTUab as features, with the exception of two basic physical characteristics (BMI and age), eight of the top 10 most important features for classifying DI were OTUs (one *Desulfovibrionales*, one *Actinomycetales*, four *Clostridiales*, one *Bacteroidales* and one *Lactobacillales*). For the model with Meta-OTUoc as the features for DI, the most important features were BMI and age, followed by two OTUs belonging to *Ruminococcaceae* and six human variables (height, latitude, elevation, the frequency of milk, alcohol, and egg intake). Interestingly, BMI and age were the most two most important features in all three models.

**Fig. 2.**
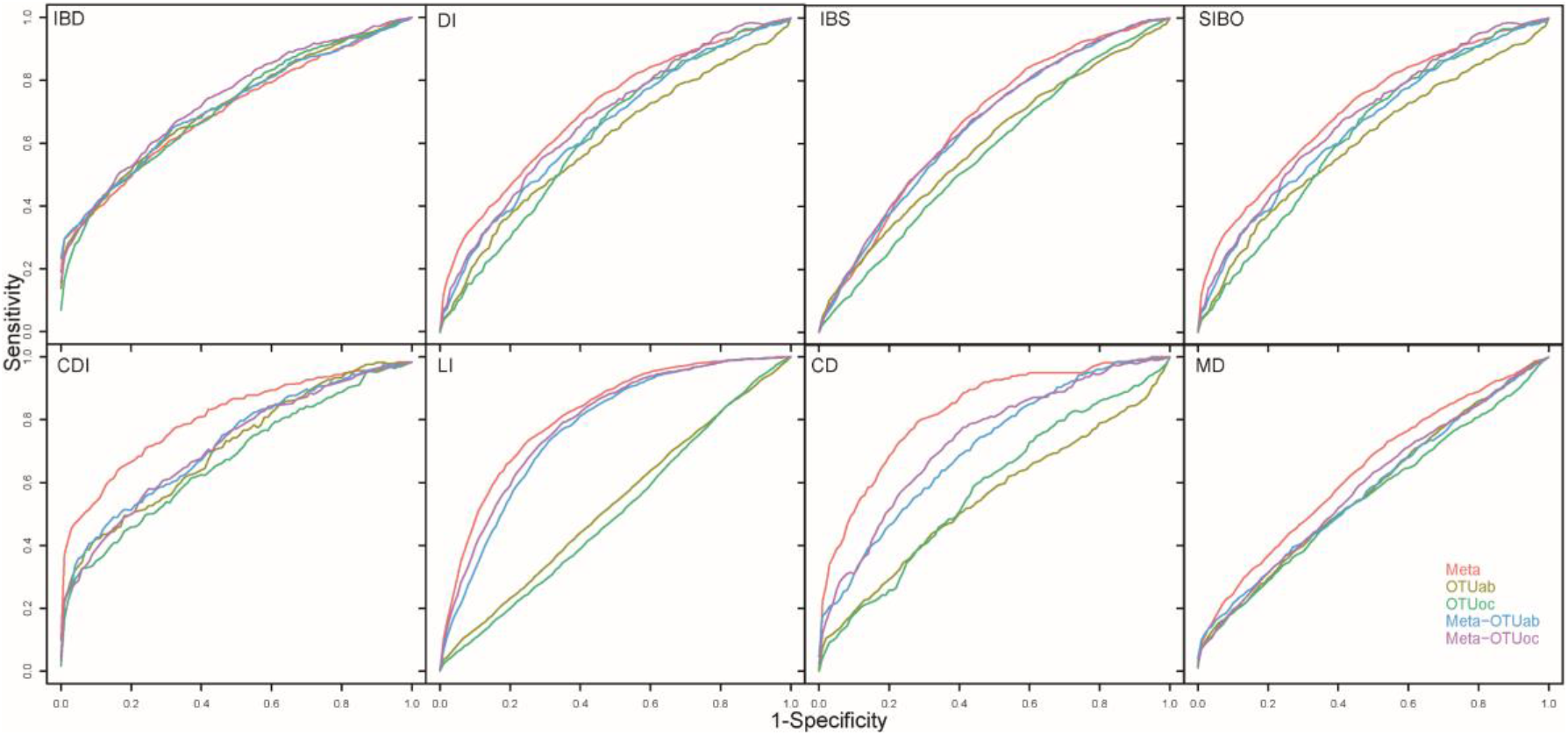
ROC curves for selection of the best classification models of eight diseases (IBD: Inflammatory Bowel Disease; CDI: C. Difficile Infection; IBS: Irritable Bowel Syndrome; SIBO: Small Intestinal Bacterial Overgrowth; DI: Diabetes; LI: Lactose Intolerance; CD: Cardiovascular Disease; MD: Mental Disorder). Line colors indicate different input features for the models.

**Fig. 3.**
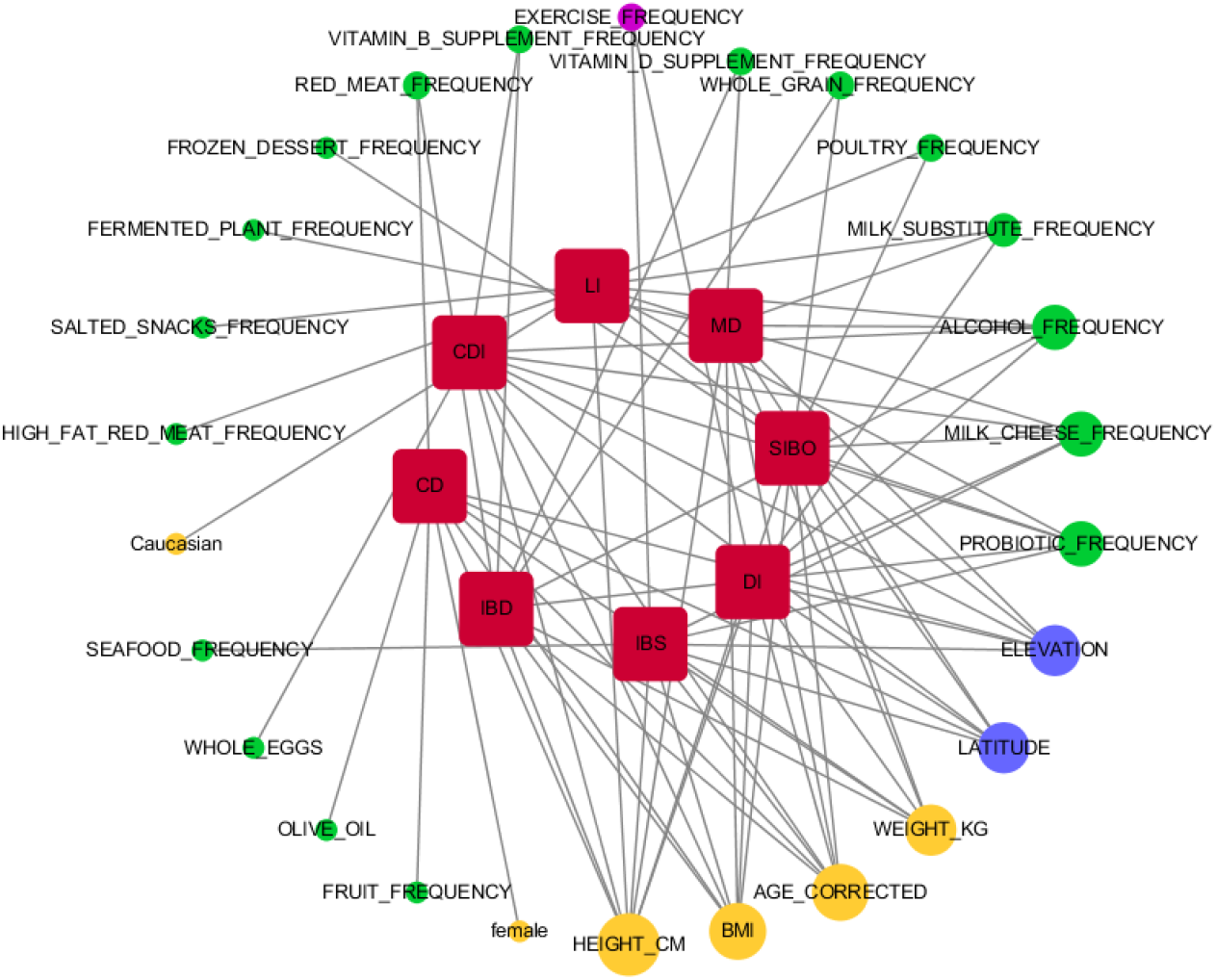
Top 10 health features for the other seven diseases (IBD: inflammatory bowel disease; CDI: C. Difficile Infection; IBS: Irritable Bowel Syndrome; SIBO: Small Intestinal Bacterial Overgrowth; DI: Diabetes; LI: Lactose Intolerance; CD: Cardiovascular Disease; MD: Mental Disorder). The direct line from one feature to one disease indicates the feature is one of the top 10 health features of the disease. The seven diseases are shown in red. The health features are shown in green, with the intensity of node shading indicating a greater disease classification value of the feature.

When classifying IBS, the best model using Meta achieved an AUC of 0.68448±0.00780. The AUCs obtained using OTUab alone (0.59820±0.00995), OTUoc alone (0.56972±0.01460) and Meta-OTUab (0.66473±0.01118) were significantly lower than those obtained using Meta. However, the AUCs obtained using Meta-OTUoc (0.66950±0.01386) did not differ significantly from those obtained using Meta alone (Meta), suggesting that adding gut microbiota occurrence features did not improve the association strength with IBS. According to the feature weights, the top 10 most important human variables for IBS using Meta and Meta-OTUoc are shown in Table S4. When using Meta as the feature set, the top 10 most important human variables consisted of two geographical location features (latitude and elevation), four basic physical characteristics (age, weight, BMI and height), three dietary characteristics (the frequencies of probiotic, cheese and seafood intake), and one lifestyle habit (exercise frequency). For the model using Meta-OTUoc as the features, with the exception of one OTU (*Enterobacter radicincitans*), the other nine of the top 10 most important features were all human variables, comprising four dietary characteristics (the frequencies of milk cheese, probiotic, whole grain, and seafood intake), one geographical location features (latitude) and four basic physical characteristics (age, weight, sex, and height).

### Adding gut microbiota reduced the association strengths with the other five diseases

Recently, gut microbes were also reported to be related to small intestinal bacterial overgrowth (SIBO), *C. difficile* infection (CDI), lactose intolerance (LI), cardiovascular disease (CD) and mental disorders (MD) (4–7, 24). As shown in Table 1, for all five diseases, the models using Meta as the features achieved the highest AUC values (SIBO: 0.74413 ± 0.0234, CDI: 0.84091 ± 0.01890, LI: 0.81647 ± 0.00751, CD: 0.83432 ± 0.01255, MD: 0.62984±0.01240). Using OTUs alone (OTUab and OTUoc) or the OTUs combined with human variables (Meta-OTUab and Meta-OTUoc) resulted in AUCs that were significantly lower than those obtained using the human variables alone (Meta). Although many gut microbes are reported to be associated with these diseases, adding gut microbiota reduced the association strengths with the other five diseases (Table 1 and Fig. 2). In addition, neither Meta nor OTUs provide good performance for classifying MD.

The top 10 most important human variables for classifying SIBO, CDI, LI, CD and MD were identified according to the feature weights (Fig. 3 and Table S5). The top 10 most important human variables for classifying SIBO comprised five dietary characteristics (the frequencies of whole grain, milk cheese, probiotic, frozen dessert and poultry intake), four basic physical characteristics (weight, BMI, height and age) and one geographical location feature (latitude). With the exception of three basic physical characteristics (height, BMI and age) and two geographical location features (latitude and elevation), five of the top 10 most important human variables for classifying CDI were related to dietary characteristics (the frequencies of probiotic, vitamin B, alcohol, whole eggs and cheese intake). It is reasonable that the two most important human variables for classifying LI are the frequencies of milk cheese and milk substitute intake and age, followed by one race-related feature (Caucasian), five dietary characteristics (the frequencies of alcohol, poultry, high-fat red meat, salted snacks and probiotic intake), height and elevation. The top 10 most important human variables for classifying CD comprised five basic physical characteristics (age, sex, height, weight and BMI), three dietary characteristics (the frequencies of high-fat red meat, olive oil and fruit intake) and two geographical location features (latitude and elevation). It is noteworthy that MD was classified mainly by four basic physical characteristics (age, BMI, height and weight), four dietary characteristics (the frequencies of milk, alcohol, vitamin D and fermented plant intake) and two geographical location features (latitude and elevation).

### Adding gut microbiota improved the association strength with unhealthy status

To investigate the potential use of the out composition to classify the health of individuals, we defined a sample as unhealthy if it was obtained from an individual with any one of the eight diseases. Finally, we obtained 2,921 unhealthy samples containing at least one of those eight diseases for training the models using OTUs or Meta alone and in combination. As shown in Table 1, the model using Meta as the features achieved an AUC of 0.70936±0.00442. The AUCs obtained using Meta-OTUab (0.75626±0.00859) and Meta-OTUoc (0.76266±0.00520) were all significantly higher than those obtained using Meta, suggesting that adding gut microbiota improved the accuracy of classifying unhealthy status. Surprisingly, the AUC obtained using OTUab (0.71976±0.00533) alone was also significantly higher than that obtained using Meta alone, indicating that unhealthy status can be classified accurately based on the abundance information of gut microbes alone. However, the AUC obtained using OTUoc (0.71976±0.00533) was not significantly different from that obtained using Meta. When calculating the weight for each feature in the models with Meta, OTUab, Meta-OTUab and Meta-OTUoc, we identified the top 10 most important features for classifying unhealthy status (Table 3). For the model with Meta as the features, the top 10 most important human variables comprised five dietary characteristics (the frequencies of cheese, probiotic, milk substitute, vitamin D and frozen dessert intake), three basic physical characteristics (age, BMI and weight) and two geographical location features (latitude and elevation). For the model using OTUab as the features, eight of the top 10 OTUs were annotated to *Clostridiales*, one to *Bacteroidales* and one to *Bifidobacteriales*. When using Meta-OTUab as the features, with the exception of four dietary characteristics (the frequencies of milk cheese, probiotic, vitamin D and milk substitute intake) and age, the other five of the top 10 most important features were all OTUs (four annotated to *Clostridiales* and one *Bacteroidales*). When using Meta-OTUoc as the features to classify unhealthy status, the top 10 most important features comprised five dietary characteristics (the frequencies of milk cheese, probiotics, vitamin D, milk substitute and frozen dessert intake), one basic physical characteristic (age), one geographical location feature (latitude) and three OTUs annotated to *Clostridiales*.

**Table 3.** Top 10 features using three types of feature sets for unhealthy status.

| Meta | OTUab | Meta-OTUab | Meta-OTUoc |
| --- | --- | --- | --- |
| MILK_CHEESE/1 | c_Clostridia;o_Clostridiales;f_Ruminococcaceae;g_Faecalibacterium;s_prausnitzii/3.2 | MILK_CHEESE/1 | MILK_CHEESE/1 |
| PROBIOTIC/2.6 | c_Clostridia;o_Clostridiales;f_;g_;s_/4.3 | PROBIOTIC/2 | PROBIOTIC/2 |
| AGE_CORRECTED/5.9 | c_Bacteroidia;o_Bacteroidales;f_Bacteroidaceae;g_Bacteroides;s_/10.3 | VITAMIN_D_SUPPLEMENT/7.1 | c_Clostridia;o_Clostridiales;f_;g_;s_/8.6 |
| MILK_SUBSTITUTE/6.4 | c_Actinobacteria;o_Bifidobacteriales;f_Bifidobacteriaceae;g_Bifidobacterium;s_adolescentis/10.4 | c_Clostridia;o_Clostridiales;f_;g_;s_/7.9 | VITAMIN_D_SUPPLEMENT/11 |
| VITAMIN_D_SUPPLEMENT/8.3 | c_Clostridia;o_Clostridiales;f_;g_;s_/11.2 | MILK_SUBSTITUTE/11.7 | MILK_SUBSTITUTE/13.1 |
| LATITUDE/9.2 | c_Clostridia;o_Clostridiales;f_Lachnospiraceae;g_Blautia;s_/14.1 | c_Clostridia;o_Clostridiales;f_Ruminococcaceae;g_Faecalibacterium;s_prausnitzii/12.5 | FROZEN_DESSERT/18.9 |
| BMI/9.3 | c_Clostridia;o_Clostridiales;f_Lachnospiraceae;g_;s_/14.7 | c_Clostridia;o_Clostridiales;f_;g_;s_/14.1 | c_Clostridia;o_Clostridiales;f_Ruminococcaceae;g_;s_/20.8 |
| ELEVATION/10 | c_Clostridia;o_Clostridiales;f_Ruminococcaceae;g_Faecalibacterium;s_prausnitzii/15.1 | AGE_CORRECTED/19.3 | c_Clostridia;o_Clostridiales;f_Ruminococcaceae;g_Oscillospira;s_/22.6 |
| FROZEN_DESSERT/10.7 | c_Clostridia;o_Clostridiales;f_Ruminococcaceae;g_;s_/22.4 | c_Clostridia;o_Clostridiales;f_;g_;s_/31 | AGE_CORRECTED/23.5 |
| WEIGHT_KG/10.7 | c_Clostridia;o_Clostridiales;f_;g_;s_/22.6 | c_Bacteroidia;o_Bacteroidales;f_Bacteroidaceae;g_Bacteroides;s_/39.8 | LATITUDE/31.5 |

## DISCUSSION

### Important OTUs for human diseases

For IBD, adding gut microbiota to human variables can achieve better results than that achieved using human variables alone and Meta-OTUoc achieved higher AUCs than that using Meta-OTUab. Among the top 10 most important features used to classify IBD with Meta-OTUoc, eight were OTUs with one belonging to the *Klebsiella* genus. *Klebsiella* is an intestinal pathobiont that can produce a cytotoxin (tillivaline), and is thought to be involved in the pathogenesis of IBD (25). It is also interesting that seven of the eight OTUs belonged to the *Clostridiales* order. At the family level, two were annotated to *Ruminococcaceae*, which have been reported as a prominent family in IBD, especially two species, *Ruminococcus torques* and *Ruminococcus gnavus* (26) and three were annotated to *Lachnospiraceae*, which is also reported to be related to IBD (27). The physiologic niche of *Ruminococcus gnavus* was speculated to be mucolytic, with dramatic changes in this species affecting the delicate equilibrium of the mucus layer and potentially increasing the intestinal permeability in IBD patients (26). For the unhealthy status classification, the AUC obtained using OTUab alone was significantly higher than that obtained using Meta alone, which shows that unhealthy status can be classified accurately using only the abundance information of gut microbes, and that gut microbes may be good biomarkers for the general health condition of humans. Eight of the top 10 OTUs with the highest weight belonged to the *Clostridiales* order. Under the *Clostridiales* order, the *Lachnospiraceae* family has been associated with many human diseases, such as inflammatory bowel disease (27), irritable bowel syndrome (28), type 1 diabetes (29, 30), *Clostridium difficile* infection (31) and liver cirrhosis (32). *Ruminococcaceae* family was also reported to be related to *Clostridium difficile* infection (31) and type 1 diabetes (29). The other two most important OTUs for unhealthy status classification belonged to the *Bacteroides* genus and the *Bifidobacterium* genus. According to the previous research, *Bacteroides* genus was associated with several human diseases, including five gut diseases (irritable bowel syndrome (28), *Clostridium difficile* infection (31), colorectal carcinoma (33), Crohn’s disease (34, 35) and infectious colitis (36)), type 1 diabetes (29, 30) and liver cirrhosis (37).

### Important human variables for human diseases

Human variables showed a strong efficiency in human disease classification. According to our results shown in Fig. 3, the basic physiological characteristics (BMI, age, height and weight) are the most important human variables correlated to most diseases, followed by location factors (latitude and elevation) and the frequency of probiotic, milk cheese and alcohol intake. BMI and age were found to be important classifiers of all seven diseases except lactose intolerance, which is supported by a cross-sectional study of pre-specified demographic and clinical data (38). Multiple pieces of evidence from experimental and observational studies showed that for a substantial proportion of patients with IBS, their symptoms were associated with the ingestion of specific foods, such as milk, which contain lactose, a disaccharide that is not effectively digested by many adults worldwide (39). Additionally, evidence exists to suggest that probiotics may exert an effect on IBS through various mechanisms (40). In accordance with previous studies, we found that milk cheese and probiotic intake were two of the most important types of human variables in addition to attitude and age for classifying IBS (Table S4). For DI classification, we found that BMI and age were the two most important types of human variables, which was supported by a cross-sectional study (38). We found that the most important health feature for classifying SIBO was the frequency of probiotic intake, which was supported by a previous systematic review (41). The review and meta-analysis showed that probiotics are both safe and effective for preventing SIBO. We also found that the frequency of cheese and milk intake are two of the most important features for LI classification. This finding is not surprising because the breakdown of nondigested lactose causes LI; therefore, LI management usually involves excluding milk and milk products from the diet. It is noteworthy that the race-related feature was also among the top 10 health features for LI classification. This discovery was validated by the previous reports that lactase persistence varies among different human populations (42). Age and sex were found to be the two most important features for classifying CD. This is a reasonable conclusion because age has been reported as one of the most powerful risk factors for developing CD (43) and there is a higher prevalence of CD in men than in women (44).

### Removing probiotics, vitamin B and vitamin D

The frequency of probiotic intake was among the top 10 features of IBD, CDI, IBS, SIBO, and LI; the frequency of eating vitamin B was among the top ten features of CDI and IBD; the frequency of vitamin D intake was among the top 10 features of IBD and MD (Fig. 3). Some samples may have been obtained from individuals who adopted these three dietary habits advised by the clinician; therefore, we repeated our analysis after removing these three health features. As the results are shown in Table 4, we found that, after removing the frequency of probiotic and vitamins B and D intake, Meta-OTUoc showed significantly better results than Meta for both classifying IBD and unhealthy status, showing that the independent associations do exist between gut microbiota and IBD and unhealthy status. It is interesting that in addition to DI and IBS, Meta-OTUoc also did not differ significantly from Meta for the classification of SIBO and MD. For the other three diseases (CDI, LI and CD), Meta still showed the best performance. The top 10 most important features for disease classification after removing the frequency of probiotic and vitamins B and D intake are shown in Table S6.

**Table 4.** Comparing the differences of AUCs of nine diseases using five feature types after removing probiotic, vitamin B and D.

| Feature type |  | Meta | OTUab | OTUoc | Meta-OT<br>Uab | Meta-OT<br>Uoc |
| --- | --- | --- | --- | --- | --- | --- |
| IBD | AUC | 0.70430±<br>0.01896 | 0.71616±<br>0.01961 | 0.70988±<br>0.01362 | 0.71616±<br>0.01731 | 0.73755±<br>0.01779 |
|  | p-value | - | 0.1578 | 0.57154 | 0.21862 | 0.00011 |
| DI | AUC | 0.85186±<br>0.01353 | 0.65737±<br>0.01907 | 0.64386±<br>0.05136 | 0.81847±<br>0.01887 | 0.83323±<br>0.02016 |
|  | p-value | - | <0.00001 | <0.00001 | 0.00406 | 0.07444 |
| IBS | AUC | 0.66737±<br>0.00874 | 0.59746±<br>0.0115 | 0.57002±<br>0.01385 | 0.65477±<br>0.00876 | 0.66202±<br>0.01747 |
|  | p-value | - | <0.00001 | <0.00001 | 0.01195 | 0.51024 |
| SIBO | AUC | 0.70714±<br>0.01773 | 0.60376±<br>0.02748 | 0.62955±<br>0.01989 | 0.65295±<br>0.02895 | 0.67997±<br>0.02833 |
|  | p-value | - | <0.00001 | <0.00001 | 0.00002 | 0.00833 |
| CDI | AUC | 0.82321±<br>0.01605 | 0.72817±<br>0.02593 | 0.68964±<br>0.01523 | 0.74306±<br>0.02055 | 0.73450±<br>0.01714 |
|  | p-value | - | <0.00001 | <0.00001 | 0.00001 | <0.00001 |
| LI | AUC | 0.81425±<br>0.00502 | 0.52895±<br>0.01525 | 0.50413±<br>0.02001 | 0.76971±<br>0.0082 | 0.78787±<br>0.00734 |
|  | p-value | - | <0.00001 | <0.00001 | <0.00001 | 0.00001 |
| CD | AUC | 0.82741±<br>0.01343 | 0.54769±<br>0.03572 | 0.57925±<br>0.01822 | 0.70966±<br>0.03159 | 0.73393±<br>0.02467 |
|  | p-value | - | <0.00001 | <0.00001 | <0.00001 | <0.00001 |
| MD | AUC | 0.63616±<br>0.01439 | 0.57518±<br>0.01564 | 0.55468±<br>0.0179 | 0.57928±<br>0.01538 | 0.58750±<br>0.02399 |
|  | p-value | - | 0.00003 | <0.00001 | 0.00016 | 0.00232 |
| UH | AUC | 0.69841±<br>0.00528 | 0.72057±<br>0.00633 | 0.71888±<br>0.0072 | 0.74820±<br>0.00376 | 0.75979±<br>0.00446 |
|  | p-value | - | 0.00007 | 0.00008 | <0.00001 | <0.00001 |

P values were calculated by comparing models' AUC scores with the models using Meta respectively and the AUC score was denoted as the mean value ± standard deviation.

### Model performance changed with OTU numbers

When there are too many OTUs as input features, the models may be overfitted. Therefore, we evaluated the changes in classification performance for the four diseases (IBD, DI, IBS and UH) in the optimal model result with the number of OTUs. The changes in AUCs obtained using the number of OTUs are shown in Fig. 4. We found that using only some of the OTUs achieved better results than using all 517 OTUs. Especially for DI, using only the top 1% of OTUs (10 OTUs) achieved the best effect. Furthermore, the top 3% of OTUs (20 OTUs) generated the best unhealthy status and IBS classification results with AUCs of 0.766 and 0.701, while the top 77% of OTUs (400 OTUs) generated the best IBD classification results with an AUC of 0.761. The OTU set with the best classification results were different for the four diseases. These OTU sets can be used as biomarkers for the corresponding diseases. In addition, when we used only gut microbes for disease classification, the OTUab achieved better results than that achieved using OTUoc, which can be explained by the greater loss of information using OTUoc than when using OTUab. However, after combination with human variables, the performance of Meta-OTUoc surpassed the Meta-OTUab in most cases. This difference might be caused by the reason that values of human variable data were associated with microbial abundance, and, for disease classification, information provided by human variables and microbial abundance were overlapped.

**Fig. 4.**
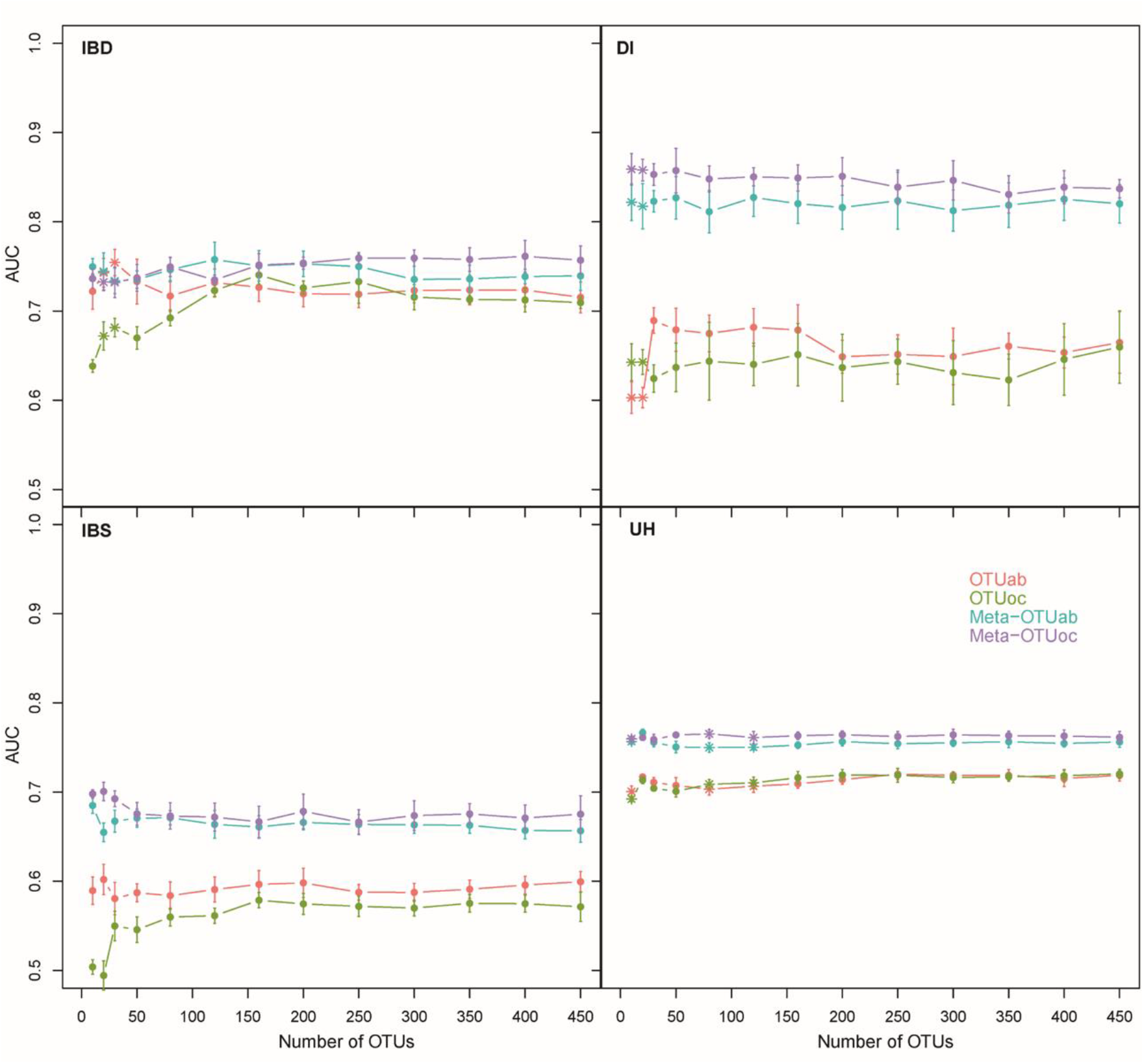
Changes in the AUC of the optimal model with the number of OTUs. The optimal model for four diseases (IBD: Inflammatory Bowel Disease; IBS: Irritable Bowel Syndrome; DI: Diabetes; UH: Unhealthy status).

## MATERIALS AND METHODS

### Data sources

We downloaded the OTU table (11-packaged/fecal/100nt/ag_fecal.biom) and human variables (11-packaged/fecal/100nt/ag_fecal.txt) from the latest version (updated in January 2018) of the AGP database available at. The original OTU table was saved as a binary file (.biom), which was converted manually to plain text with Python Script, which is available at GitHub (https://github.com/tinglab/kLDM.git). The original gut microbial abundance table (OTU table) contained 15,158 samples and 24,114 OTUs selected by applying a 97% similarity cutoff with SortMeRNA defined by the AGP consortium. The OTUs were mapped to the Greengenes Database (45) to identify their taxonomy. Each cell in the OTU table presents the abundance of its corresponding OTUs in a specific sample. The original human variables file (Meta table) contained 15,158 samples and 523 factors related to physicochemical parameters of fecal samples, dietary habits, lifestyle choices and some diseases. Each cell in the Meta table presents the measured value of its corresponding meta-data in a specific sample.

### Data preprocessing

We selected 30 items of human variables to classify disease, including individuals’ physiological characteristics, lifestyle, location, and diet. Among these 30 items, six were related to physiological characteristics; two were associated with lifestyle choices; three were associated with location; the remaining 19 were connected with diet (frequencies of fermented plant, frozen dessert, fruit, high-fat red meat, home-cooked meals, alcohol, red meat, meat eggs, milk substitute, milk cheese, olive oil, probiotic, salted snacks, seafood, vegetable, vitamin D supplement, vitamin B supplement, whole grain and whole eggs). The values of frequency-related human variables were categorized as follows: ‘Never’, ‘Rarely’ (less than once/week), ‘Occasionally’ (1–2 times/week), ‘Regularly’ (3–5 times/week) and ‘Daily’. For convenience, these categories were recoded as integers from 1 to 5 (where 1 represents ‘never’ and 5 represents ‘daily’) according to their frequencies (Table S1). The values of human variables were missing for some samples; therefore, the samples with complete sets of 30 human variable data items were selected for the following analysis. Samples with huge (first 1%) and small (last 2%) reads, as well as those with evenness <2 were removed. For all selected samples, OTUs were filtered based on their average size and non-zero times. Finally, 7,571 samples with 517 OTUs and complete human variables were obtained.

Eight diseases (CD, SIBO, MD, LI, DI, IBD, IBS, CDI and DI) that have been reported in previous studies to be related to gut microbiome were selected. The samples from individuals affected by any of these eight diseases were treated as unhealthy (UH). Information about individuals’ disease status was extracted, and the samples from individuals with diseases were labeled. The characteristics of the dataset, the demographic details of samples and the number of male and female patients for each disease are shown in Table S2.

### Machine learning models training and evaluation

To evaluate the confound association between disease with gut microbiota and human variables, four machine learning (ML) techniques (RF, Random Forest; GBDT, Gradient Boosting Decision Tree; LR, Logistic Regression; XGBoost, eXtreme Gradient Boosting) were used to build the model, and the AUC scores were calculated to compare their performance (Fig. 1). For every disease, four types of ML models were trained with five-fold cross-validation using training data, including 90% of all samples, and the model providing the best performance was selected based on the maximal AUC. Considering that the positive samples labeled with diseases occupied a tiny proportion, an equal number of negative samples were randomly selected for model training. The optimal model was then evaluated and compared using the validation dataset comprising the remaining 10% of samples.

In addition to the different model types, five combinations of features were used to construct the following separate models to capture the best features for classifying each disease: human variables only (Meta), OTU abundance only (OTUab), OTU occurrence only (OTUoc), both human variable data and OTU abundance (Meta-OTUab), and both human variable data and OTU occurrence (Meta-OTUoc). The OTU occurrence was determined based on the existence of OTUs only. Models using different combinations of features were trained and compared using an identical dataset. For each type of feature, the best model with the maximal AUC score was selected from the four types of models.

To assess the significance of differences in the model performance among the five types of features, the model training process was repeated 10 times with a random selection of training data. The AUC scores were presented as the mean±standard deviation. For each disease, paired sample *t*-tests were used to compare the differences in AUC values between the feature type ‘Meta’ and the four other feature types (’OTUab’, ‘OTUoc’, ‘Meta-OTUab’ and ‘Meta-OTUoc’). In the statistical analysis, Bonferroni correction was used to adjust for the multiple testing error. In considering nine disease and four comparisons, we tested 36 independent hypotheses using the same data at the 0.05 significance level, and instead of using a *P*-value threshold of 0.05, we use a stricter threshold of 0.0014.

### Identification of microbial biomarkers of diseases

For the best performing model of each disease, the weights of each feature were calculated. The top ten features (OTUs or Meta) with the highest absolute weights were selected as the biomarkers for the disease. We then obtained the taxa of these OTUs and verified their relationships to the disease by searching published literature or databases. In this study, we used the Human Microbe-Disease Association Database (HMDAD) (http://www.cuilab.cn/hmdad), which is a curated collection of microbe-disease association from previous microbiota studies. The OTUs with high weights that were verified were treated as microbial biomarkers of the disease.

## ACKNOWLEDGMENT

This research was supported by the National Natural Science Foundation of China (Grant NO. 61872218 and 61721003), China Postdoctoral Science Foundation (Grant NO. 2020M680454), Tsinghua University Students Research Training (Grant NO. 53412002520) and Beijing National Research Center for Information Science and Technology (BNRist). The funders had no role in study design, data collection and analysis, decision to publish, or preparation of the manuscript.

All authors conceived the study and participated in its design. CMZ, XW performed the collecting and cleaning of datasets. XW, YQY analyzed the data and performed the model analysis. CMZ, JCL, RJ and TC performed manuscript preparation. All authors read and approved the final manuscript.

## Supplementary Tables

**Table S1.**
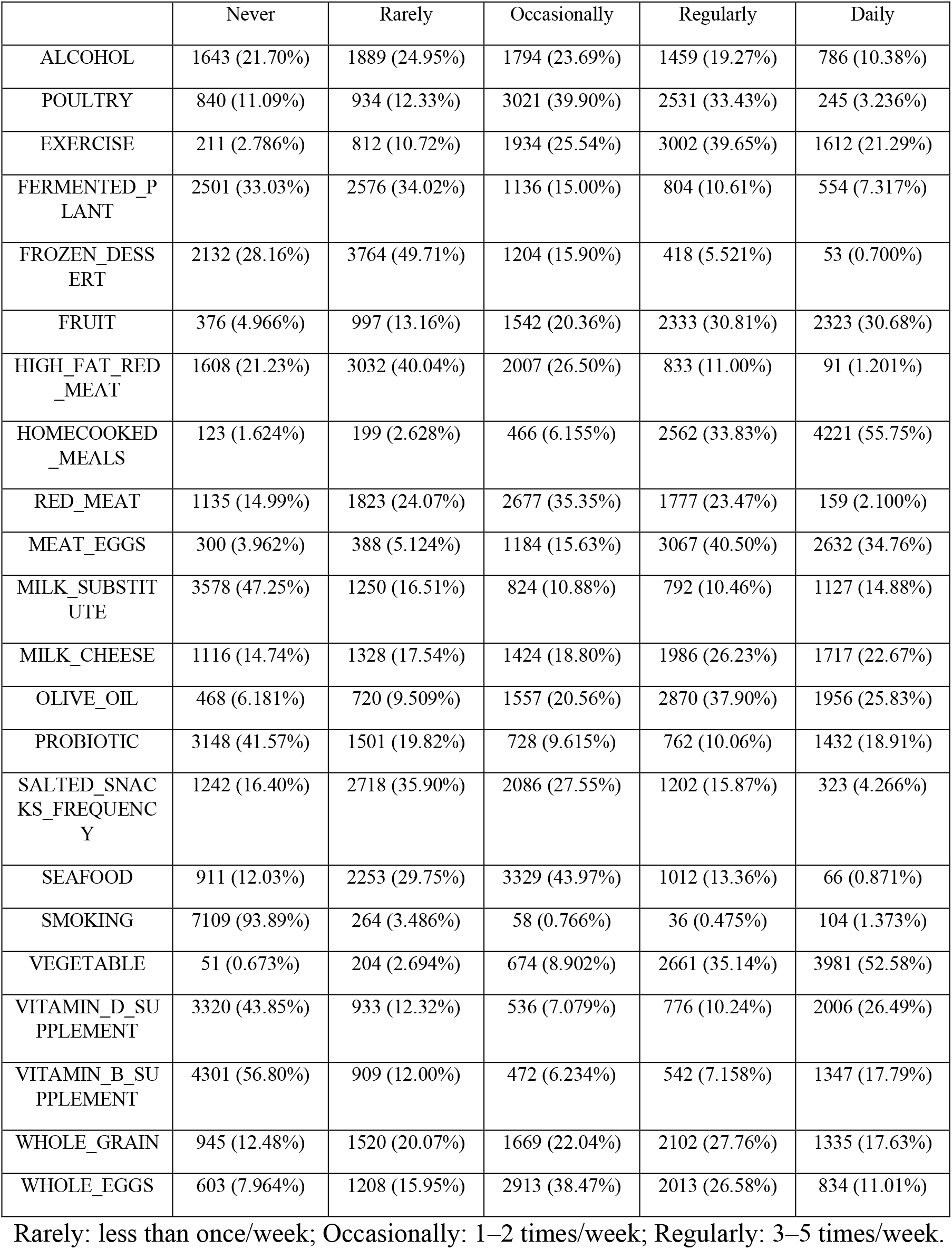
Frequency of diet and lifestyle factors.

**Table S2.** Basic information of the dataset.

|  | Female | Male | Total |
| --- | --- | --- | --- |
| Number | 3983 | 3588 | 7571 |
| Age | 47(35,59) | 46(33,61) | 47(34,59) |
| BMI | 22.43(20.36,25.63) | 24.28(22.05,26.61) | 23.4(20.98, 26.3) |
| Height (cm) | 165(160,170) | 178(172,183) | 170(162,178) |
| Weight (kg) | 61(55,70) | 78(69,87) | 69(58,81) |
| Latitude | 40.8(34.1,51.3) | 40.1(33,50.175) | 40.7(33.6,51) |
| Elevation | 81.2(28.4,193.3) | 96.1(30.9,205.375) | 88.1(29.8,200.6) |
| Caucasian | 3575 | 3186 | 6761 |
| Asian or Pacific Islander | 162 | 178 | 340 |
| African American | 29 | 16 | 45 |
| Hispanic | 97 | 102 | 199 |
| Other Race | 120 | 106 | 226 |
| CARD | 74 | 180 | 254 |
| SIBO | 209 | 101 | 310 |
| MENTAL | 407 | 270 | 677 |
| LACTOSE | 712 | 508 | 1220 |
| IBS | 666 | 333 | 999 |
| IBD | 164 | 249 | 413 |
| Diabetes | 74 | 100 | 174 |
| CDIFF | 92 | 87 | 179 |
| Unhealth | 1612 | 1309 | 2921 |

The representation format of the values for age, BMI, height, weight, latitude and elevation are median (lower quartile and upper quartile).

**Table S3.**
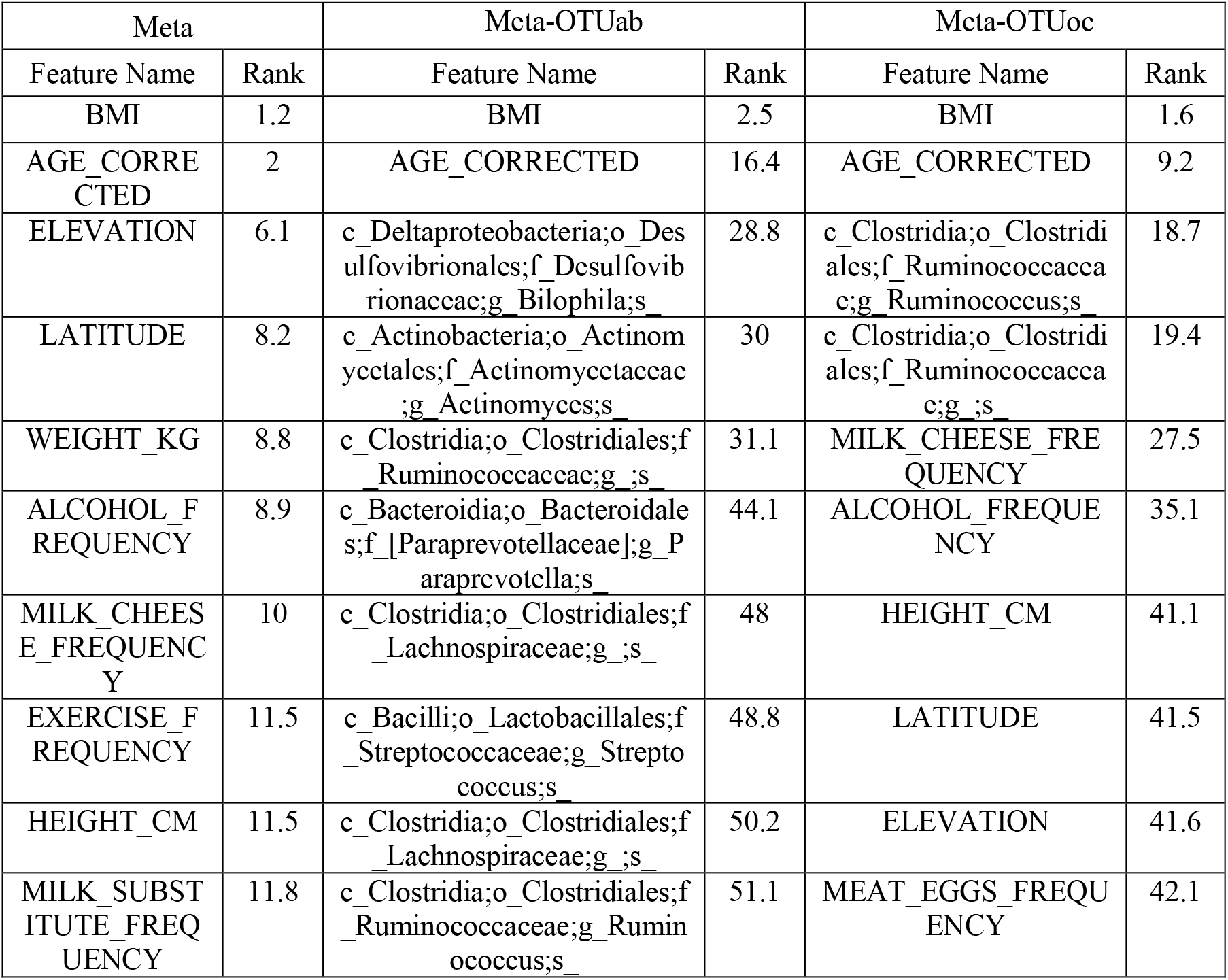
Top 10 features using three types of feature sets for Diabetes.

**Table S4.** Top 10 features using two types of feature sets for IBS.

| Meta |  | Meta-OTUoc |  |
| --- | --- | --- | --- |
| Feature Name | Rank | Feature Name | Rank |
| LATITUDE | 1.8 | MILK_CHEESE | 3.8 |
| AGE_CORRECTED | 2.5 | PROBIOTIC | 4 |
| PROBIOTIC | 4.6 | LATITUDE | 4.9 |
| MILK_CHEESE | 5 | AGE_CORRECTED | 8.7 |
| WEIGHT_KG | 6.6 | WEIGHT_KG | 10.7 |
| ELEVATION | 6.8 | c_Gammaproteobacteria;o_Enterobacteriales;f_Enterobacteriaceae;g_Enterobacter;s_radicincitans | 17 |
| BMI | 7.1 | female | 26.9 |
| EXERCISE | 10.7 | HEIGHT_CM | 28 |
| HEIGHT_CM | 11 | WHOLE_GRAIN | 28.8 |
| SEAFOOD | 11.5 | SEAFOOD | 30.3 |

**Table S5.**
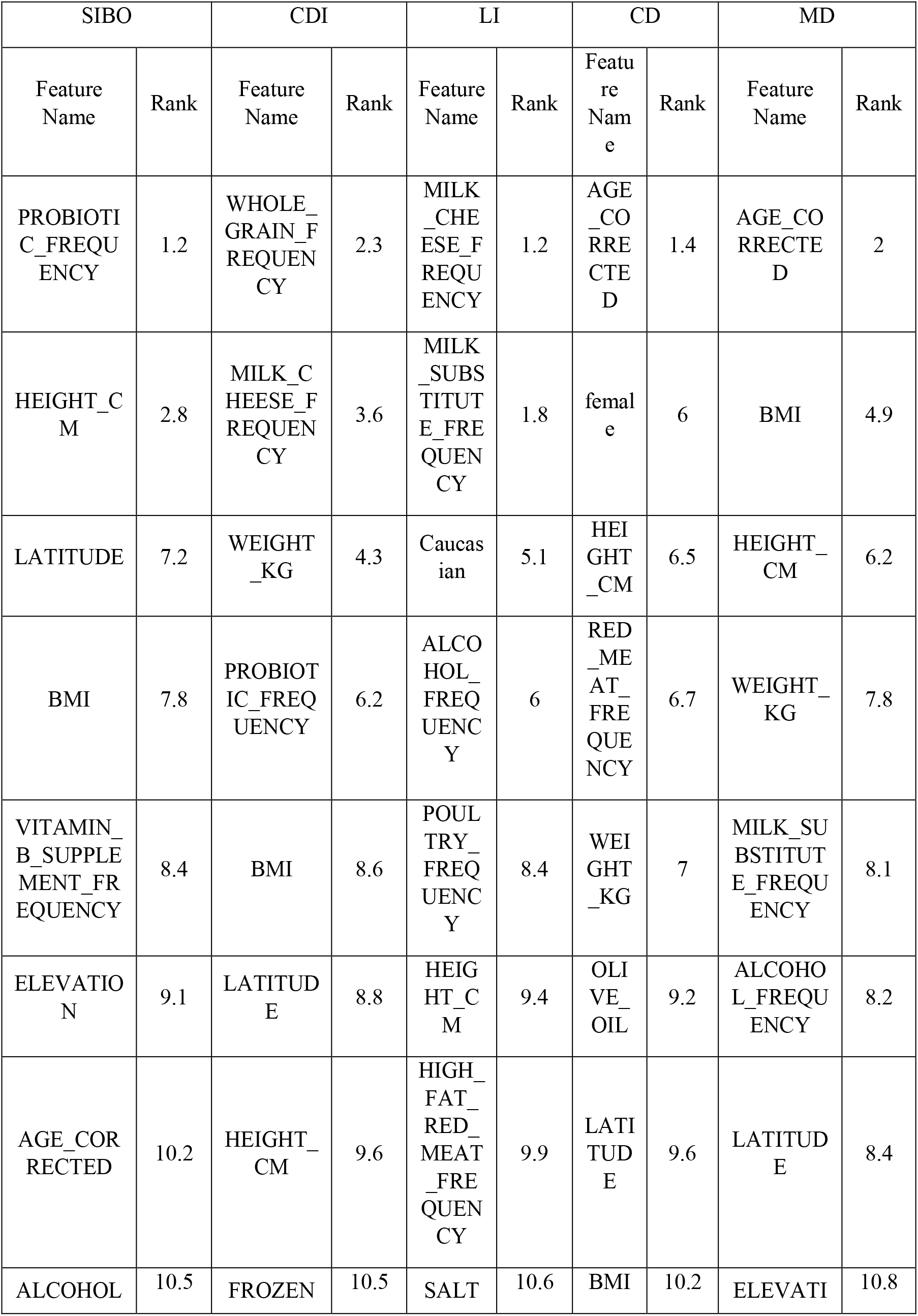
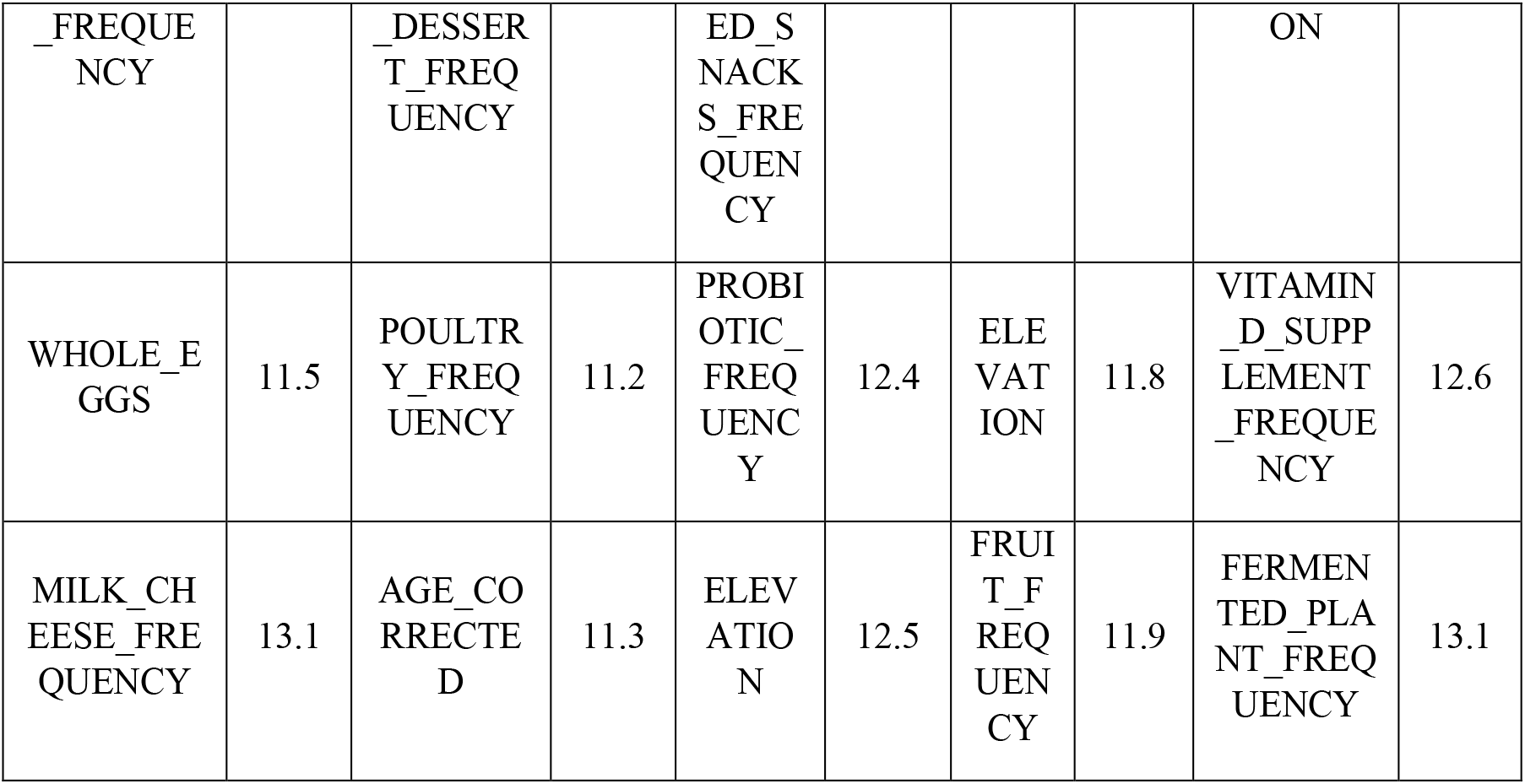
Top 10 features using human variables for the other five diseases.

**Table S6.**
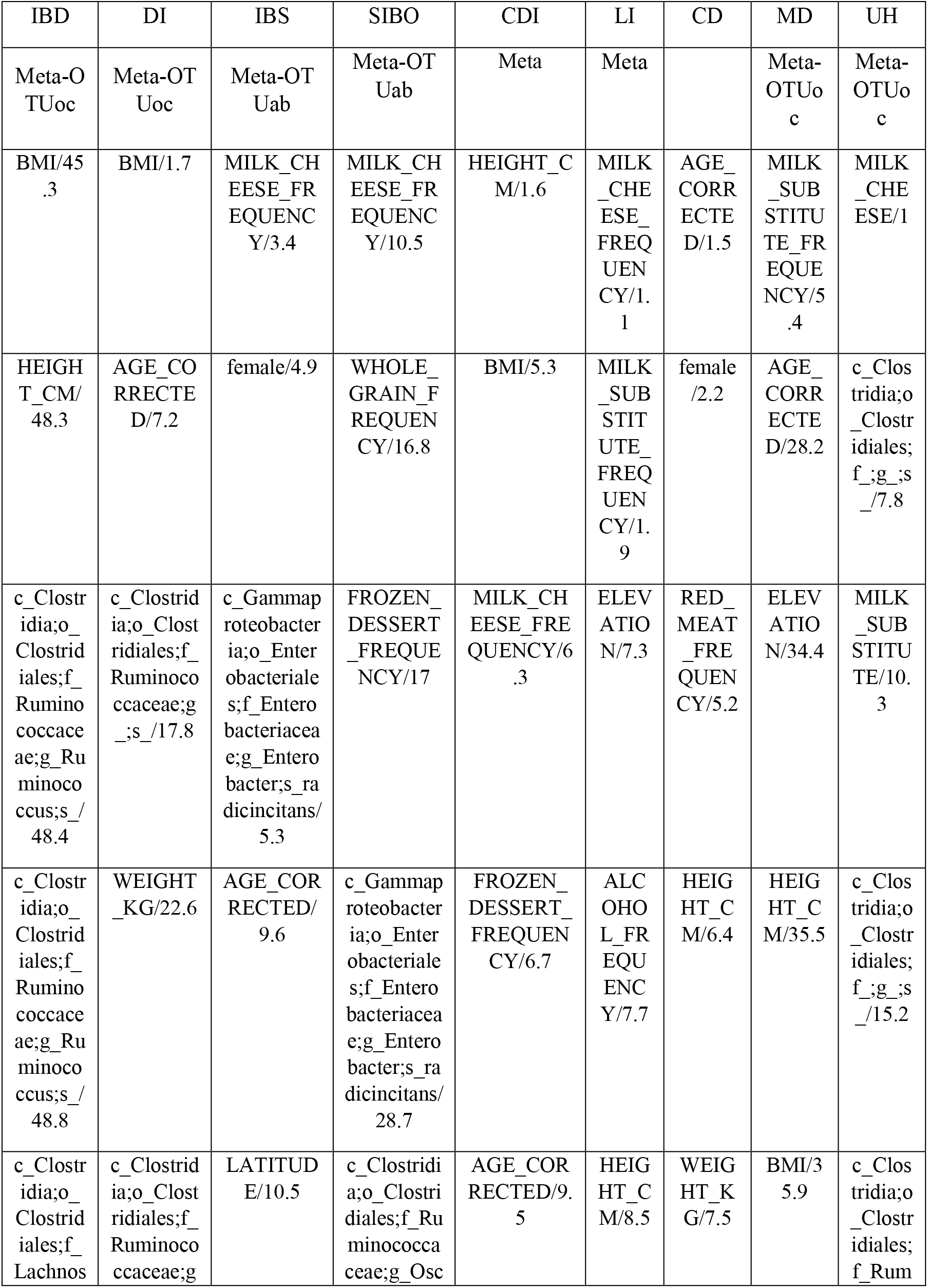
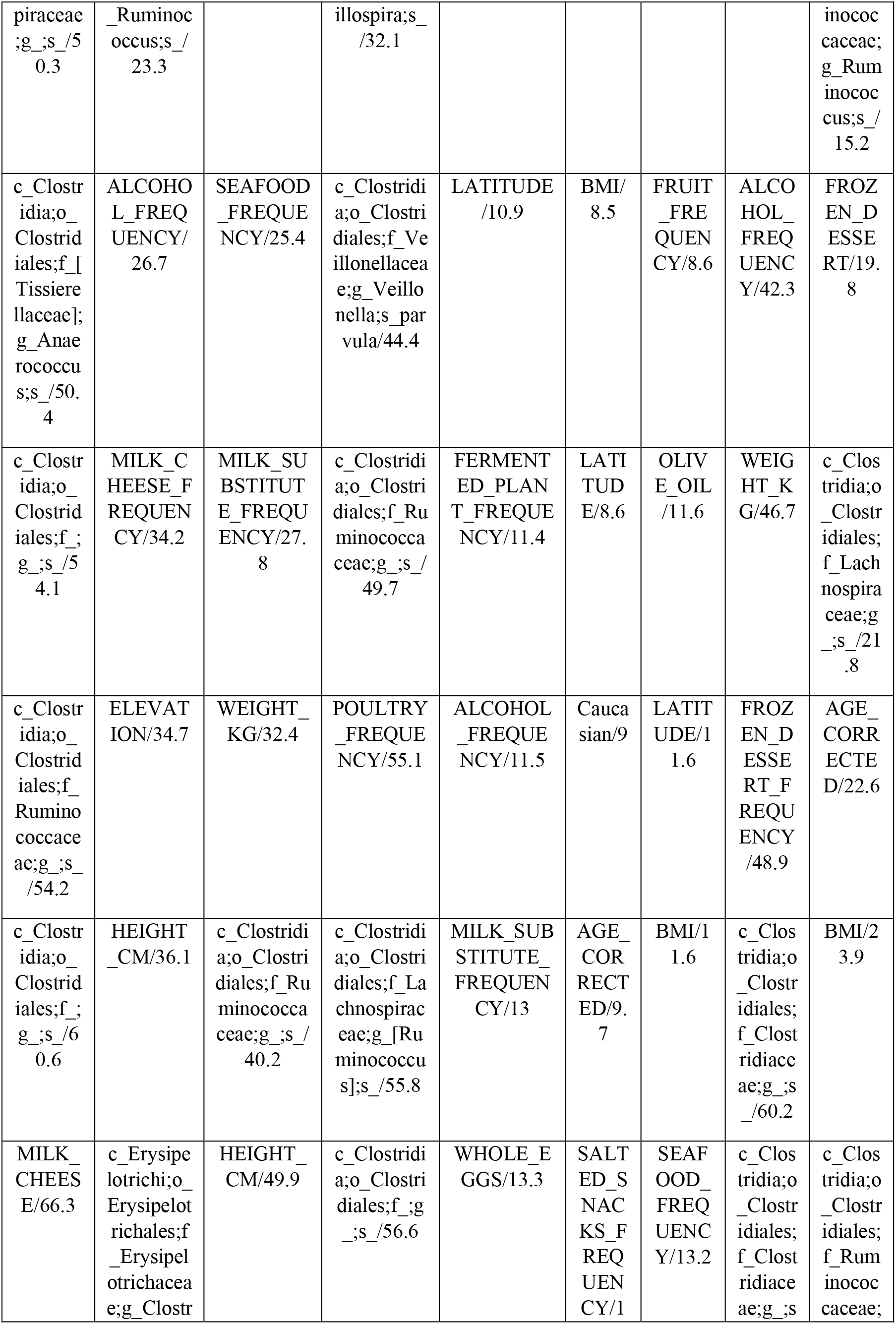
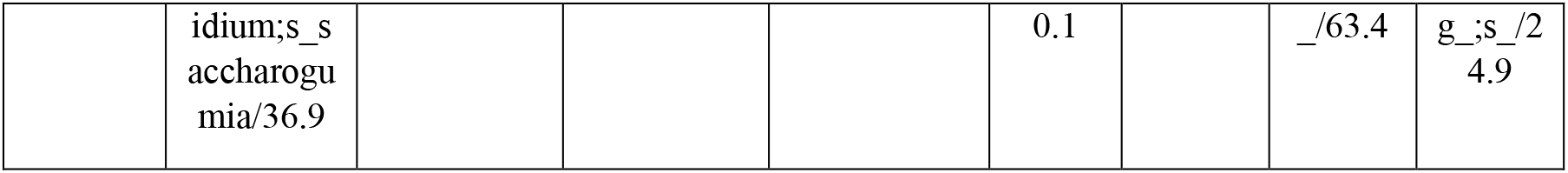
Top 10 features after removing probiotics, vitamin B and vitamin D for all diseases.

